# Dose-Dependent Effects of Low-Intensity Focused Ultrasound on Human Deep White Matter Tracts

**DOI:** 10.64898/2026.09.02.748985

**Authors:** Aki Tsuchiyagaito, Masaya Misaki, Xi Ren, Danielle Clark, Adrienne Taren, Chieh V. Chen, Maria Ironside, Mirta F. Villarreal, Rayus Kuplicki, Sanjay Mathew, Martin Paulus, Noah S. Philip, Salvador M. Guinjoan

## Abstract

**Background:** Low-intensity focused ultrasound (LIFU) is emerging as a method for anatomically specific, noninvasive, and reversible neuromodulation of deep brain structures relevant to psychopathology. A key requirement for clinical translation is evidence of graded target engagement. We examined whether LIFU applied to deep white matter tracts produces dose-dependent functional connectivity changes in healthy adults.

**Methods:** In this preregistered, double-blind, randomized study, 35 healthy adults underwent LIFU targeting (80 s, 10% duty cycle, estimated ISPPA 2.26 W/cm^2^, 0.5 MHz, in a θ-burst pattern) of individually modeled right thalamo-prefrontal white matter tracts under two dose conditions: one and three consecutive sonications. Dose-dependent changes in resting-state functional connectivity were examined during a 30-minute poststimulation period in 31 participants.

**Results:** One sonication acutely increased right thalamic functional connectivity with the left thalamus (28 voxels; Z = 5.48, Cohen’s d = 0.61), whereas three sonications decreased connectivity with the right dorsolateral prefrontal cortex (Z = -4.05, Cohen’s d = -0.45). Poststimulation time-course analysis revealed a 3-LIFU-specific linear recovery trend in the left cerebellum (Z = 4.14, d = 0.48), reflecting initial suppression followed by progressive recovery across 30 minutes. Stronger thalamo-prefrontal structural connectivity was associated with condition-specific functional connectivity changes (left frontal pole; F = 14.73, ηp² = 0.41).

**Conclusions:** These findings provide initial human evidence that LIFU is well tolerated and produces dose-dependent changes in functional connectivity associated with deep white matter tract stimulation, informing parameter selection for future circuit-based mechanistic and clinical studies.

## Introduction

Internalizing disorders, including major depressive disorder (MDD) and post-traumatic stress disorder, are leading psychiatric causes of disability and death by suicide (1). Whereas about two-thirds of patients with MDD respond to first-line treatments such as medications and psychotherapy, a substantial proportion do not respond and are considered treatment-resistant (2). The last two decades have witnessed intense research efforts to address this problem by directly targeting dysfunctional circuits, both invasively and noninvasively. The most established noninvasive approach to treating these disorders is repetitive transcranial magnetic stimulation (TMS) (3), which is advancing from empirical to hypothesis-based targeting strategies designed to modify specific, anatomically circumscribed circuits, thereby maximizing efficacy and limiting side effects (4–6). Although TMS is transforming noninvasive neuromodulation, its direct biophysical reach is largely constrained to superficial cortical tissue. Deeper limbic targets implicated in affective psychopathology must therefore generally be engaged indirectly through connected cortical regions, as in depression protocols that stimulate superficial sites connected with the subgenual cortex rather than the structure itself (7–9). H-coils can modulate deeper structures, but without comparable anatomical specificity (10). The ability to focally modulate deep structures and their interconnecting white matter pathways is highly relevant because internalizing symptoms are increasingly understood as manifestations of distributed cortico-subcortical circuit dysfunction rather than isolated regional abnormalities (11–17). Accordingly, most neurosurgical approaches to refractory internalizing syndromes converge on deep white matter connections (18), including capsulotomy (19), deep brain stimulation (DBS) of the anterior limb of the internal capsule (20), and cingulotomy (21). Even DBS of the subgenual cingulate (22) is understood to derive its beneficial effects from the large-scale connectivity of white matter tracts underlying the initially identified Brodmann area 25 cortical target, rather than from this gray matter region alone (23). In this context, in vitro evidence indicates that myelinated axon function is particularly sensitive to disruption by low-intensity focused ultrasound (LIFU) (24–27). This provides a mechanistic rationale for using LIFU for reversible, anatomically specific, and noninvasive modulation of deep myelinated axon bundles, with broad potential clinical applications. Preclinical studies demonstrate that LIFU alters saltatory conduction through effects on TRAAK and TREK potassium channels located at nodes of Ranvier (28). These channels are highly sensitive to mechanical and thermal energy (27), and their opening can produce axonal hyperpolarization through increased K^+^ efflux, thereby attenuating saltatory conduction (28). We recently reported proof-of-concept evidence supporting translation of these preclinical observations to patients with major depression, in whom functional MRI connectivity, heart rate variability, and behavioral measures indicated anatomically specific deep white matter target engagement (29). Demonstrating dose-dependent effects is a critical next step because it would establish graded, measurable changes in the intended neural system and support anatomically specific biological target engagement rather than nonspecific effects (30,31). It would also help define a therapeutic window: stimulation parameters sufficient to modulate the relevant circuit while remaining safe, tolerable, and anatomically precise. This provides the mechanistic and safety rationale needed before clinical efficacy trials. We therefore conducted a preregistered, double-blind, randomized study of two LIFU doses applied to thalamo-prefrontal white matter tracts in healthy individuals (NCT07099950). We hypothesized that the intensity (study aim 1) and duration (study aim 3) of LIFU-related changes in resting-state functional connectivity of the thalamus would differ by dose. We also hypothesized that neuromodulation intensity would be directly related to the structural connectivity of the targeted axon bundles, which provide the anatomical pathway for putative functional connectivity changes (study aim 2).

## Methods and Materials

### Participants and design

Thirty-five healthy adults were enrolled between April 22, 2025, and April 30, 2026, in a preregistered, randomized, double-blind crossover study (NCT07099950). The study was approved by the Western Institutional Review Board, and all participants provided written informed consent. Supplemental Table 1 presents the inclusion and exclusion criteria, and Table 1 summarizes participant demographic characteristics. A CONSORT-style diagram in the Supplement describes participant flow. Figure 1, panel A, depicts the general study design. Each participant completed three visits: a baseline visit (V0) without LIFU stimulation and two LIFU visits (Visits 1 and 2) in randomized order. Participants received one LIFU epoch (1-LIFU) at one visit and three consecutive epochs (3-LIFU) at the other (Figure 1, panel B). At all visits, participants underwent MRI scanning as described below.

**Figure 1.**
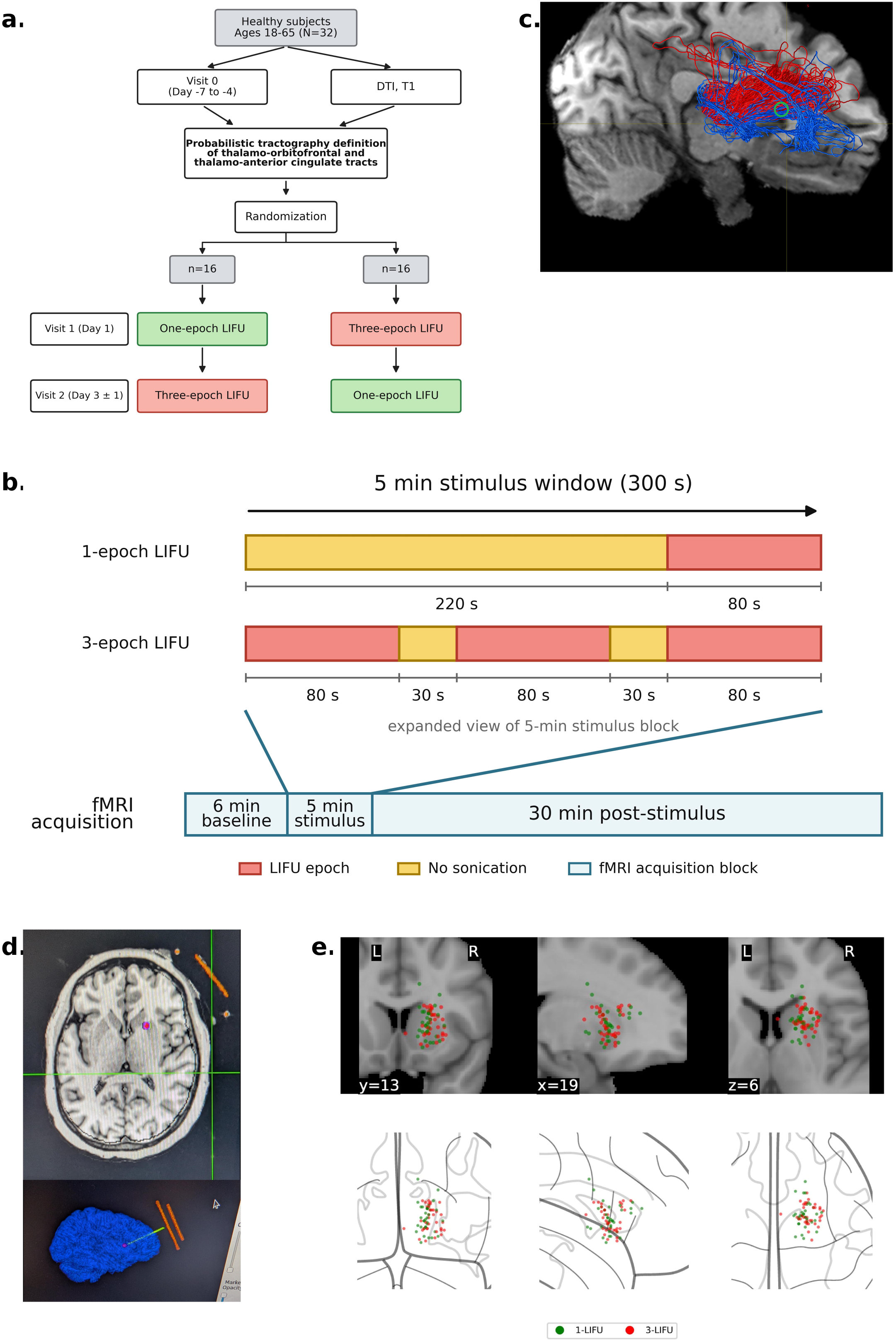
Study design and targeting. (a) General study design. (b) One-epoch and three-epoch LIFU conditions within the continuous 41-min resting-state fMRI acquisition. Red bars indicate LIFU sonication epochs, and yellow bars indicate intervals without sonication. (c) Example probabilistic tractography streamlines connecting the right thalamus with the anterior cingulate cortex (red) and orbitofrontal cortex (blue); the green circle marks the selected target. (d) Custom software used to model the ultrasound beam and define the sonication target in the scanner. (e) Final sonication targets for the 1-LIFU (green) and 3-LIFU (red) conditions. DTI, diffusion tensor imaging; fMRI, functional magnetic resonance imaging; LIFU, low-intensity focused ultrasound.

**Table 1.**
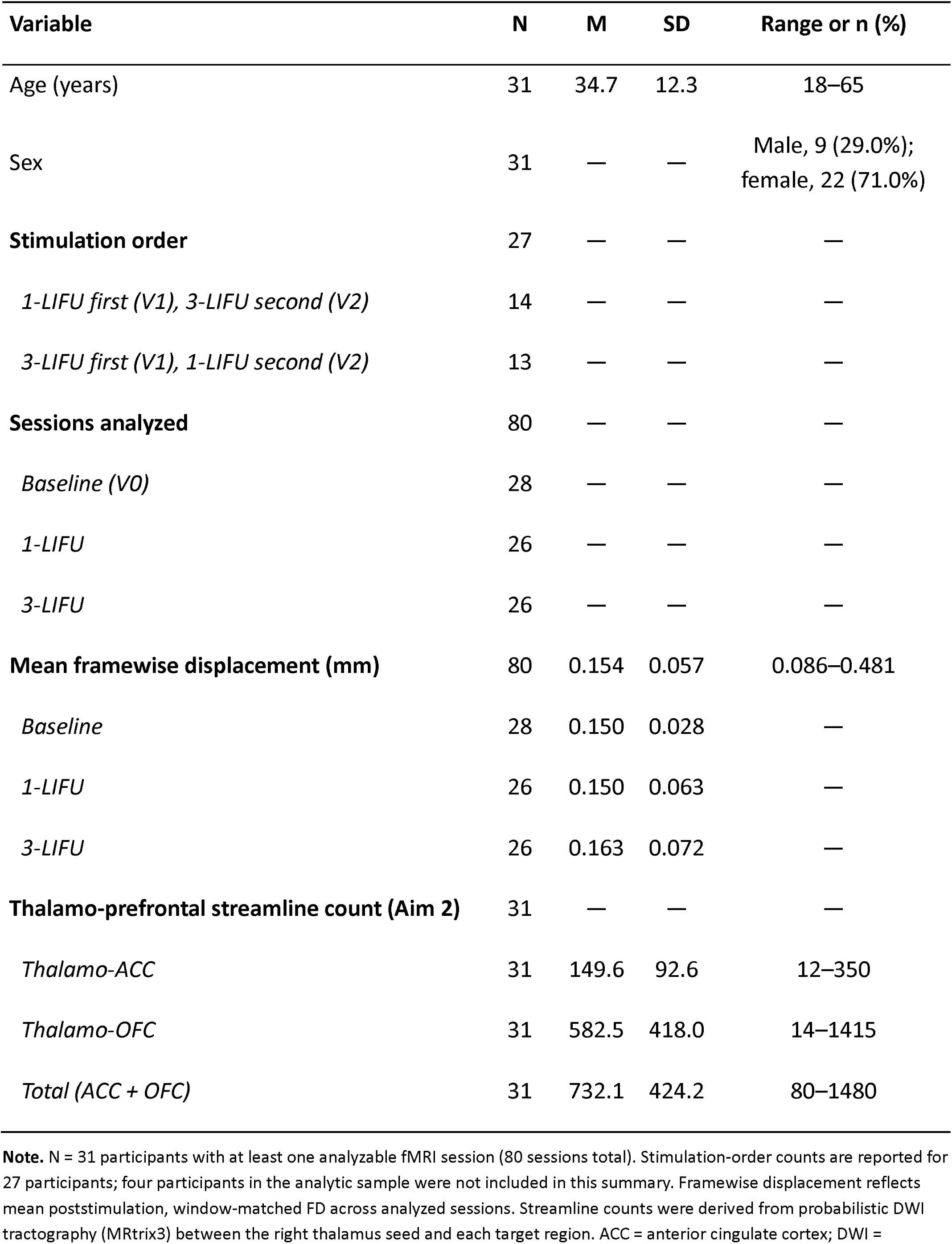
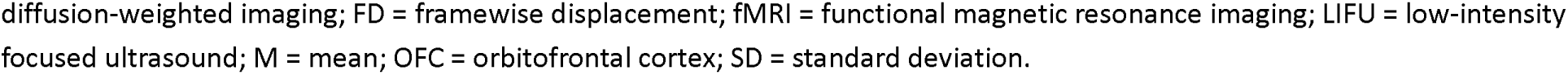
Participant and Analyzed-Session Characteristics (N = 31)

| **Variable** | **N** | **M** | **SD** | **Range or n (%)** |
| --- | --- | --- | --- | --- |
| Age (years) | 31 | 34.7 | 12.3 | 18–65 |
| Sex | 31 | — | — | Male, 9 (29.0%);<br>female, 22 (71.0%) |
| **Stimulation order** | 27 | — | — | — |
| *1-LIFU first (V1), 3-LIFU second (V2)* | 14 | — | — | — |
| *3-LIFU first (V1), 1-LIFU second (V2)* | 13 | — | — | — |
| **Sessions analyzed** | 80 | — | — | — |
| *Baseline (V0)* | 28 | — | — | — |
| *1-LIFU* | 26 | — | — | — |
| *3-LIFU* | 26 | — | — | — |
| **Mean framewise displacement (mm)** | 80 | 0.154 | 0.057 | 0.086–0.481 |
| *Baseline* | 28 | 0.150 | 0.028 | — |
| *1-LIFU* | 26 | 0.150 | 0.063 | — |
| *3-LIFU* | 26 | 0.163 | 0.072 | — |
| **Thalamo-prefrontal streamline count (Aim 2)** | 31 | — | — | — |
| *Thalamo-ACC* | 31 | 149.6 | 92.6 | 12–350 |
| *Thalamo-OFC* | 31 | 582.5 | 418.0 | 14–1415 |
| *Total (ACC + OFC)* | 31 | 732.1 | 424.2 | 80–1480 |

**Note.** N = 31 participants with at least one analyzable fMRI session (80 sessions total). Stimulation-order counts are reported for 27 participants; four participants in the analytic sample were not included in this summary. Framewise displacement reflects mean poststimulation, window-matched FD across analyzed sessions. Streamline counts were derived from probabilistic DWI tractography (MRtrix3) between the right thalamus seed and each target region. ACC = anterior cingulate cortex; DWI =
diffusion-weighted imaging; FD = framewise displacement; fMRI = functional magnetic resonance imaging; LIFU = low-intensity focused ultrasound; M = mean; OFC = orbitofrontal cortex; SD = standard deviation.

### LIFU stimulation

Low-intensity focused ultrasound was applied to individually modeled thalamo-prefrontal white matter tracts. Probabilistic tractography defined white matter pathways connecting the right thalamus with the anterior cingulate cortex (ACC) and orbitofrontal cortex (OFC), as described previously (32) (Figure 1, panel C). Custom software was used to define the sonication target within the scanner environment (Figure 1, panel D). We used the same stimulation parameters described elsewhere (29,32). Briefly, stimulation consisted of a θ-burst pattern of 400 20-ms pulses delivered every 180 ms, representing a 10% duty cycle and a total stimulation duration of 80 s (32). The free-field spatial-peak pulse-average intensity (I_SPPA_) was 9.06 W/cm^2^, producing an estimated tissue-level intensity of 2.26 W/cm^2^. Consistent with the US Food and Drug Administration’s nonsignificant risk determination, delivered energy was not adjusted across participants for cranial thickness. As in the previous pilot study, fixed parameters were used to minimize the risk of excessive energy delivery from participant-specific programming errors (29). The 1-LIFU condition consisted of one stimulation epoch, whereas the 3-LIFU condition consisted of three consecutive epochs separated by two 30-s intervals. The total stimulus-delivery period in the scanner was 5 min (Figure 1, panel B). Blinding was assessed immediately after each session by asking participants whether they believed they had received one or three LIFU epochs and to rate their confidence on a 0-10 scale.

### MRI acquisition and preprocessing

At baseline (V0), MRI acquisition included T1-weighted anatomical imaging, a single 6-min resting-state fMRI time series, and diffusion tensor imaging (DTI). Figure 1, panel B, shows the schema for each experimental visit (Visits 1 and 2). Each visit consisted of a continuous 41-min resting-state scan comprising a 6-min prestimulation period, a 5-min stimulation period, and a 30-min poststimulation period. The assigned LIFU condition was delivered during the stimulation period. Transducer localization was confirmed on a prestimulation image using custom software (Figure 1, panel D). The same software controlled stimulus delivery while maintaining blinding of participants, MRI operators, and study staff. In the 1-LIFU condition, one 80-s sonication was administered at the end of the 5-min stimulation period. In the 3-LIFU condition, three 80-s sonications were separated by 30-s intervals without stimulation (Figure 1, panel B). Resting-state fMRI data were preprocessed using fMRIPrep v25.2.5 with MNI152NLin2009cAsym 2-mm standard-space output and a sessionwise anatomical reference. Three participants and 15 sessions were excluded because of excessive head motion (more than 50% of volumes censored), and one participant dropped out after the first experimental visit, leaving 80 sessions from 31 participants for analysis. Nuisance regression was performed in AFNI v25.2.16 using a 12-parameter motion model (six motion parameters and their temporal derivatives), five white matter and five cerebrospinal fluid aCompCor components, and 13 physiological regressors derived from respiratory and cardiac signals (five respiratory-volume-per-time regressors and eight RETROICOR regressors). Data were bandpass filtered (0.01-0.1 Hz) and spatially smoothed within the brain mask using a 4-mm full-width-at-half-maximum kernel. Three dummy volumes were discarded. Volumes exceeding a framewise displacement threshold of 0.3 mm or a standardized derivative of root mean square variance threshold of 1.5 were censored; one TR immediately before and one immediately after each flagged volume were also censored.

#### Structural connectivity analysis

(Figure 1, panel C). Diffusion-weighted imaging data were processed using MRtrix3 as previously described (29,33) to generate probabilistic tractography streamlines between a right thalamus seed and each ipsilateral target region. Figure 1, panel C, shows an exemplar, with streamlines connecting to the ACC in red and those connecting to the OFC in blue (29). Streamline counts for the thalamo-ACC and thalamo-OFC pathways were extracted for each participant and used as continuous moderator variables. The correlation between ACC and OFC streamline counts was r = -0.04, indicating minimal correlation between the two structural connectivity measures. The tractography analysis was also used to select each participant’s sonication target, which was kept constant across experimental visits. As described elsewhere (29), two investigators (AT and SMG) independently identified the coordinates of maximum streamline density connecting to both cortical destinations. The independently selected coordinates were within 5 mm in 90% of cases. The final experimental coordinate was determined by consensus, usually at the midpoint between the two coordinates (29).

#### Functional connectivity analysis

Pearson correlations were computed between the mean time series of the right thalamus seed region and every other brain voxel and then Fisher z-transformed. Poststimulation resting-state data were divided into five consecutive 6-min windows, yielding a 30-min poststimulation time course. The single 6-min V0 resting-state scan served as the common baseline reference for comparisons with each poststimulation window from Visits 1 and 2. Additional details of the functional connectivity analysis are provided in the Supplement.

### Statistical analysis

Group-level analyses were conducted using linear mixed-effects models (AFNI 3dLME) with subject-specific random intercepts and Type III sums of squares. All models included age, sex, and window-matched mean framewise displacement as covariates. Three preregistered primary analyses were performed. Aim 1 modeled functional connectivity as a function of condition (baseline, the initial 6-min post-1-LIFU window, and the initial 6-min post-3-LIFU window), with pairwise contrasts between conditions. Aim 2 tested whether total thalamo-prefrontal structural connectivity (defined as the sum of tractography streamlines connecting the right thalamus with the right ACC and right OFC) moderated stimulus-related functional connectivity changes. Aim 3 tested whether the temporal profile of functional connectivity changes across the 30-min recovery period differed by condition, modeling the condition-by-time-window interaction with time window treated as a categorical factor and including post hoc linear-trend and decay contrasts. An exploratory analysis tested whether within-visit functional connectivity change from pre-LIFU to poststimulation was dose dependent (V1/V2 sessions only). Whole-brain cluster correction was applied using 3dClustSim (autocorrelation function parameters: a = 0.611, b = 4.016, c = 2.180; NN2 connectivity; two-sided testing), with a voxel-level threshold of p < .001 and a cluster-level FDR corrected threshold of p < .05, corresponding to a minimum cluster size of 25 voxels within the whole-brain mask (268,341 voxels).

## Results

Table 1 summarizes the demographic characteristics of the sample. Figure 1, panel E, shows the final sonication targets defined using the custom software (Figure 1, panel D) for the 1-LIFU (green) and 3-LIFU (red) conditions. One participant was excluded before targeting because faulty diffusion-weighted image acquisition prevented accurate definition of the probabilistic tractography streamlines.

Figure 2 and Table 2 summarize functional connectivity effects during the initial 6-min poststimulation period. The 1-LIFU condition produced a significant increase in right thalamic functional connectivity relative to baseline in the contralateral thalamus (mediodorsal/prefrontal nucleus; MNI peak: 3, 23, 10; Z = 5.48, Cohen’s d = 0.61; Figure 2, panel A). The 3-LIFU condition produced a significant decrease in right thalamic functional connectivity relative to baseline in the ipsilateral dorsolateral prefrontal cortex (superior/middle frontal gyrus, BA9/46; MNI peak: 27, 41, 35; Z = -4.05, Cohen’s d = -0.45; Figure 2, panel B). No clusters survived correction for the direct 3-LIFU versus 1-LIFU comparison.

**Figure 2.**
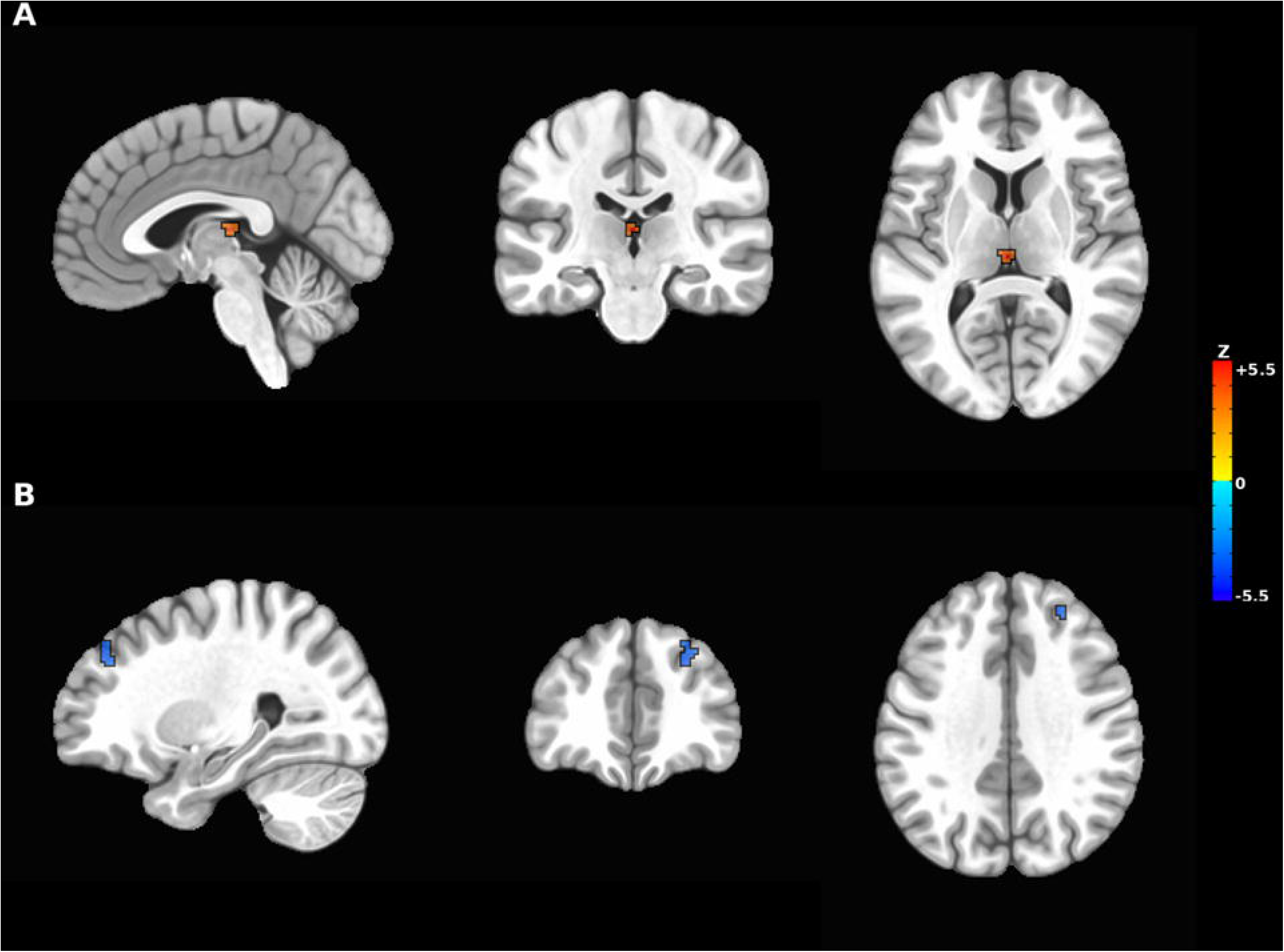
Functional connectivity effects during the initial 6-min poststimulation period. (A) The 1-LIFU condition increased functional connectivity between the right thalamus and contralateral thalamus. (B) The 3-LIFU condition decreased functional connectivity between the right thalamus and ipsilateral superior/middle frontal gyrus, a destination of the sonicated white matter tracts. Warm colors indicate positive Z statistics, and cool colors indicate negative Z statistics. Maps are displayed at voxel p < .001 and cluster-corrected p < .05, with a minimum cluster size of 25 voxels. fMRI, functional magnetic resonance imaging; LIFU, low-intensity focused ultrasound.

**Table 2.** Aim 1: Clusters of Significant Difference in Thalamic Functional Connectivity (Acute Poststimulation Window, 0-6 min)

| Region | k | x | y | z | Z | d |
| --- | --- | --- | --- | --- | --- | --- |
| *MNI coordinates (peak)* |  |  |  |  |  |  |
| **1-LIFU > Baseline** |  |  |  |  |  |  |
| L Thalamus<br>(mediodorsal/prefrontal nucleus) | 28 | 3 | 23 | 10 | 5.48 | 0.61 |
| **Baseline > 3-LIFU** |  |  |  |  |  |  |
| R Dorsolateral Prefrontal Cortex<br>(SFG/MFG, BA9/46) | 31 | 27 | 41 | 35 | -4.05 | -0.45 |
| **3-LIFU vs. 1-LIFU** |  |  |  |  |  |  |
| *No clusters survived correction* |  |  |  |  |  |  |

**Note.** k = cluster size in voxels (2 mm isotropic). MNI coordinates indicate the peak voxel. Z = peak Z statistic. d = Cohen's d ( $Z/\sqrt{N}$ , $N = 80$ sessions). Cluster correction: voxel $p < .001$ , cluster FDR-corrected $p < .05$ , minimum $k = 25$ voxels (3dClustSim, NN2, two-sided, whole-brain mask). Positive Z indicates greater functional connectivity relative to baseline; negative Z indicates reduced connectivity.

Figure 3 and Table 3 depict the progression of LIFU effects across conditions over the 30-min poststimulation period. The overall condition-by-time interaction was significant in the left superior and middle temporal gyri (BA22; 30 voxels; MNI peak: -55, -19, -7; F = 4.35, Fcrit = 3.36, ηp² = 0.09; Figure 3, panel A). A 3-LIFU-specific linear recovery trend was observed in the left cerebellum (lobules VIII and IX; 25 voxels; MNI peak: -17, -57, -47; Z = 4.14, Cohen’s d = 0.48; Figure 3, panel B), reflecting initial suppression during the first 6-min poststimulation window followed by progressive recovery over the remainder of the 30-min period. Decay contrasts did not survive cluster correction. At an uncorrected p < .001, 3-LIFU showed numerically greater suppression than 1-LIFU during the 6- to 18-min poststimulation period. In the plotted trajectories, 1-LIFU showed increased connectivity that moved toward baseline over the same period.

**Figure 3.**
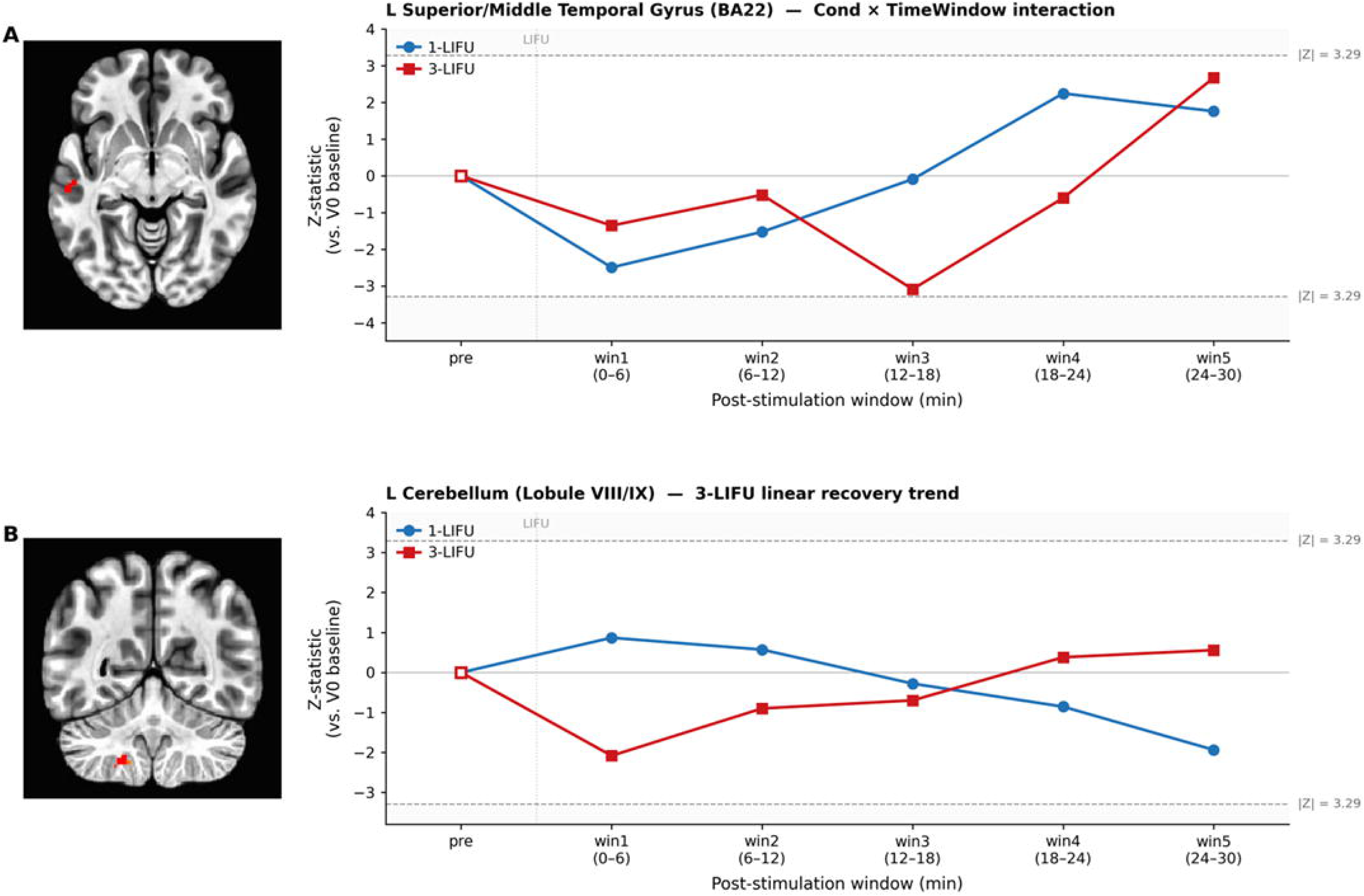
Temporal profiles of functional connectivity after 1-LIFU and 3-LIFU. Lines show model-estimated Z statistics relative to the V0 baseline across five consecutive 6-min poststimulation windows. Blue circles indicate 1-LIFU, and red squares indicate 3-LIFU. Dashed horizontal lines indicate |Z| = 3.29, corresponding to a two-sided voxel threshold of p = .001. (A) Condition-by-time-window interaction in the left superior/middle temporal gyrus (BA22). (B) 3-LIFU-specific linear recovery trend in the left cerebellum (lobules VIII/IX). BA, Brodmann area; LIFU, low-intensity focused ultrasound.

**Table 3.**
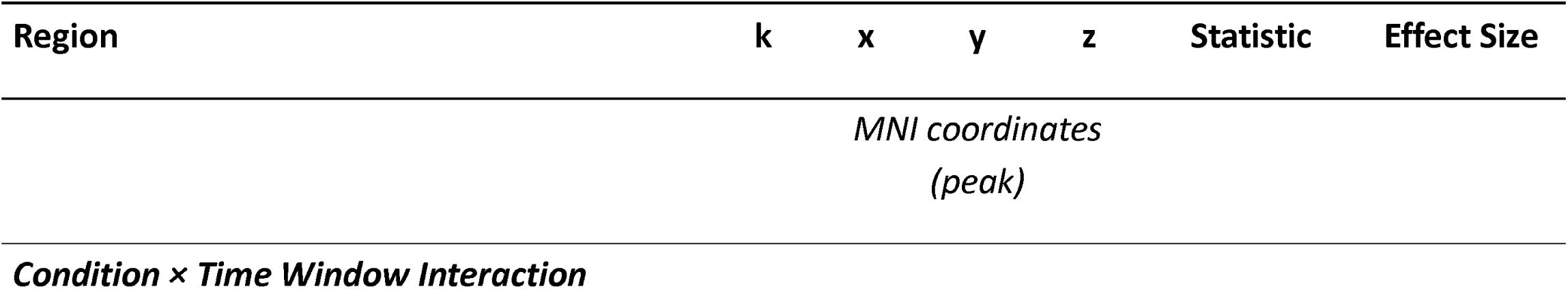
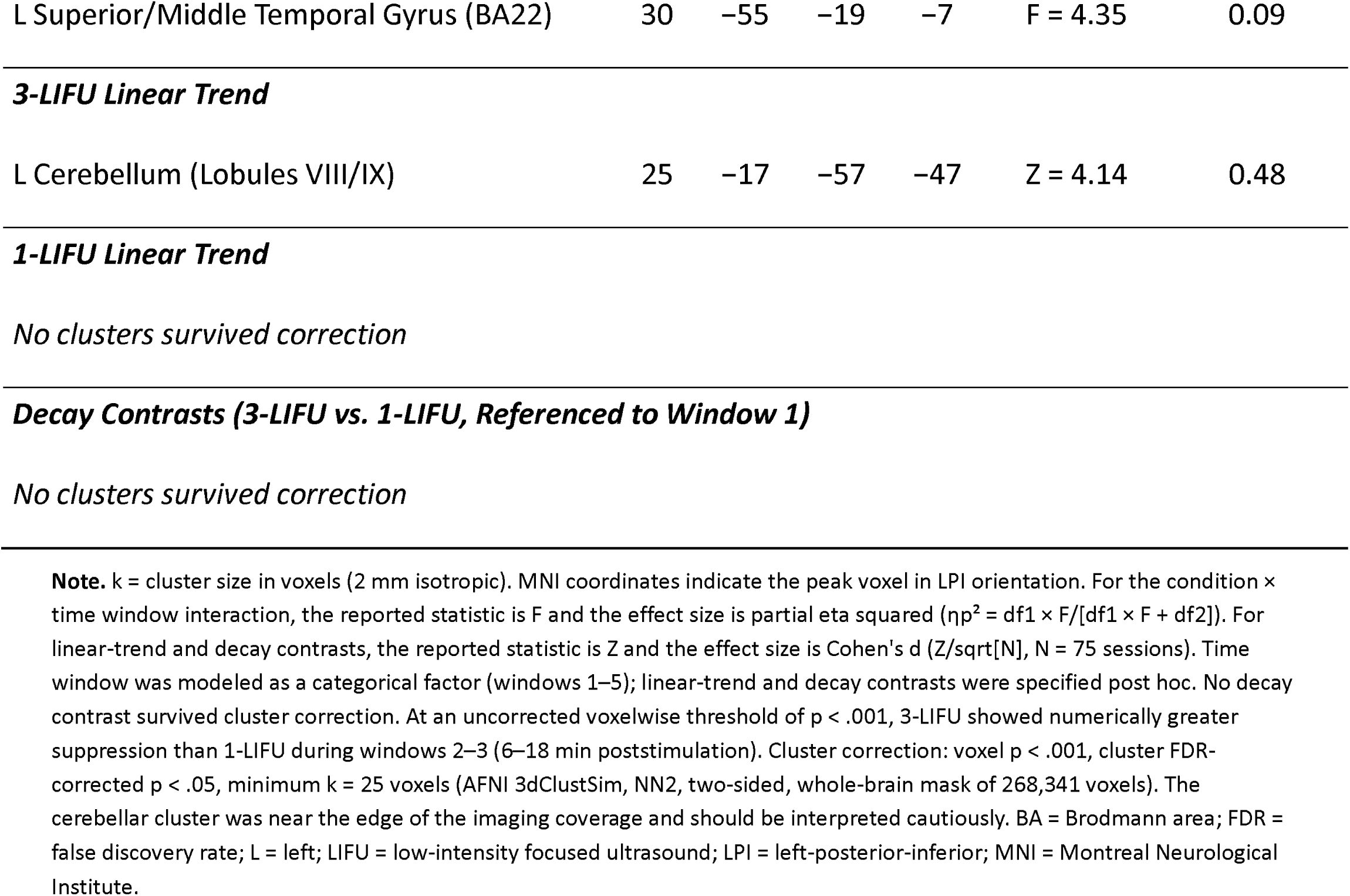
Aim 3: Clusters Showing a Significant Condition × Time Window Interaction and Post Hoc Contrasts for the Temporal Profile of Thalamic Functional Connectivity (30-Min Poststimulation Period)

| Region | k | x | y | z | Statistic | Effect Size |
| --- | --- | --- | --- | --- | --- | --- |
| *MNI coordinates (peak)* |  |  |  |  |  |  |
| **Condition × Time Window Interaction** |  |  |  |  |  |  |
| L Superior/Middle Temporal Gyrus (BA22) | 30 | -55 | -19 | -7 | F = 4.35 | 0.09 |
| 3-LIFU Linear Trend |  |  |  |  |  |  |
| L Cerebellum (Lobules VIII/IX) | 25 | -17 | -57 | -47 | Z = 4.14 | 0.48 |

Figure 4 and Table 4 summarize the relationship between total thalamo-prefrontal structural connectivity, measured as the number of probabilistic tractography streamlines connecting the right thalamus with the right OFC and ACC, and sonication-related functional connectivity changes. Greater total streamline count (ACC + OFC combined; n = 31) was significantly associated with functional connectivity changes across stimulation conditions (condition × streamline-count interaction: F = 17.13, df1 = 2, df2 = 43). The omnibus interaction produced significant clusters in the left paracentral lobule/supplementary motor area (57 voxels; MNI peak: -33, -27, 66; F = 17.13, ηp² = 0.44), left frontal pole (BA10; 49 voxels; MNI peak: -9, -77, -11; F = 14.73, ηp² = 0.41), and right fusiform/inferior temporal gyrus (BA37; 34 voxels; MNI peak: 46, 36, -13; F = 13.71, ηp² = 0.39) (Figure 4, panel A). Post hoc condition contrasts are shown in Figure 4, panels B and C. For 1-LIFU relative to baseline, greater streamline count was associated with increased thalamic functional connectivity in the left thalamus (29 voxels; MNI peak: -3, -23, 10; Z = 5.42, d = 0.61) and right inferior temporal gyrus/temporal pole (BA20/38; 29 voxels; MNI peak: 52, 8, -31; Z = 5.42, d = 0.61) (Figure 4, panel B). For 3-LIFU relative to baseline, greater streamline count was associated with decreased functional connectivity between the right thalamus and ipsilateral dorsolateral prefrontal cortex (BA9/46; 32 voxels; MNI peak: 28, 42, 36; Z = -4.77, d = -0.54) (Figure 4, panel C).

**Figure 4.**
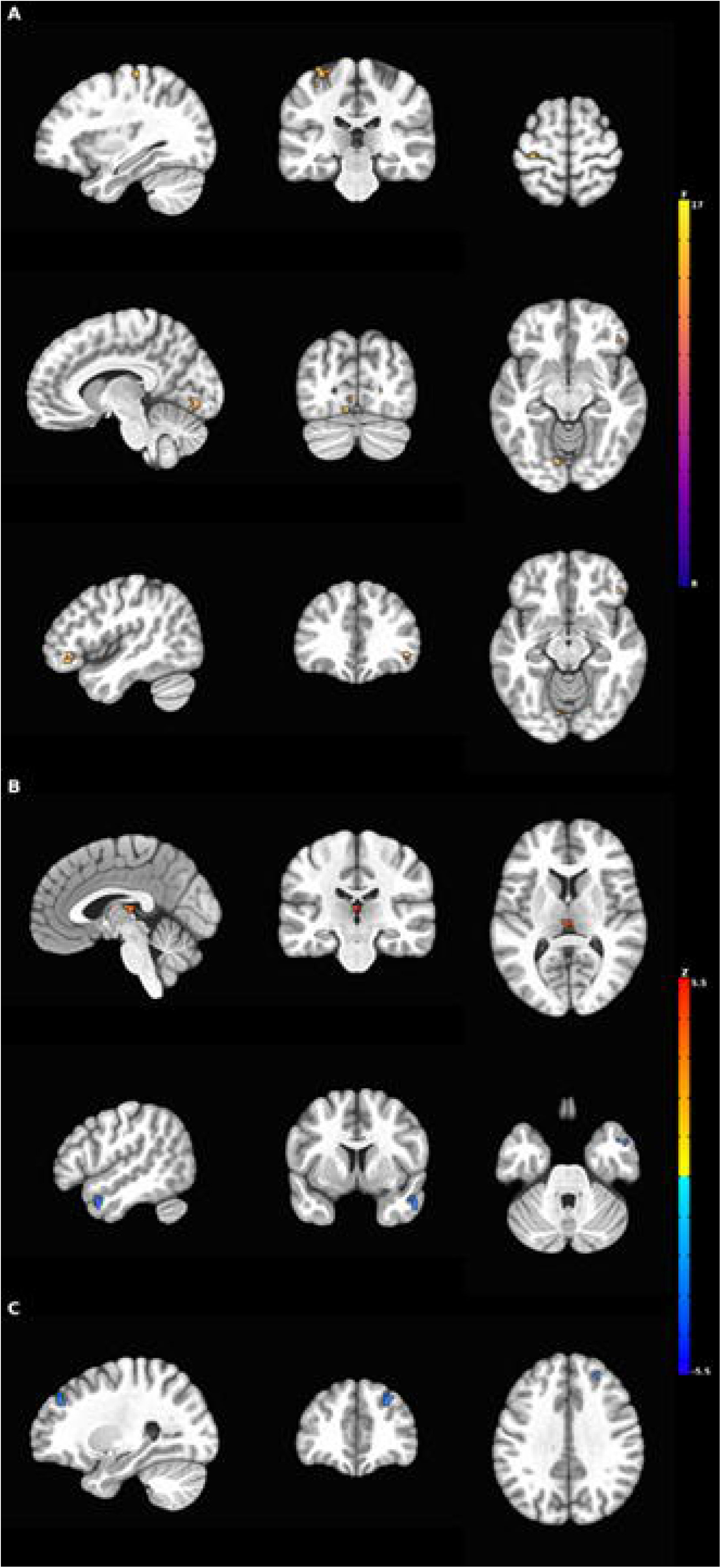
Association between thalamo-prefrontal structural connectivity and poststimulation functional connectivity changes. (A) Omnibus condition × streamline-count interaction, with clusters in the left paracentral lobule/supplementary motor area, left frontal pole, and right fusiform/inferior temporal gyrus. (B) For 1-LIFU relative to baseline, greater streamline count was associated with increased functional connectivity in the left thalamus and right inferior temporal gyrus/temporal pole. (C) For 3-LIFU relative to baseline, greater streamline count was associated with decreased functional connectivity in the right dorsolateral prefrontal cortex. Panel A displays F statistics; panels B and C display Z statistics. Warm colors indicate positive associations, and cool colors indicate negative associations. Maps are displayed at voxel p < .001 and cluster-corrected p < .05, with a minimum cluster size of 25 voxels. LIFU, low-intensity focused ultrasound.

**Table 4.**
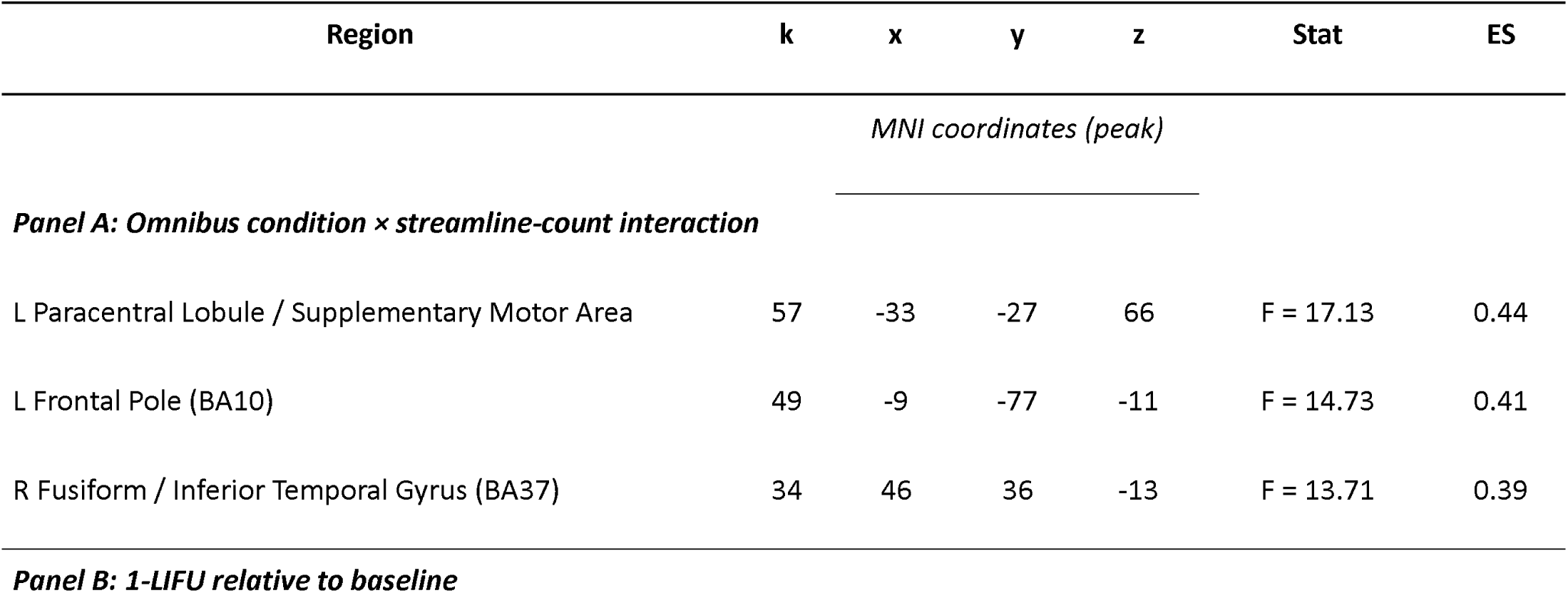
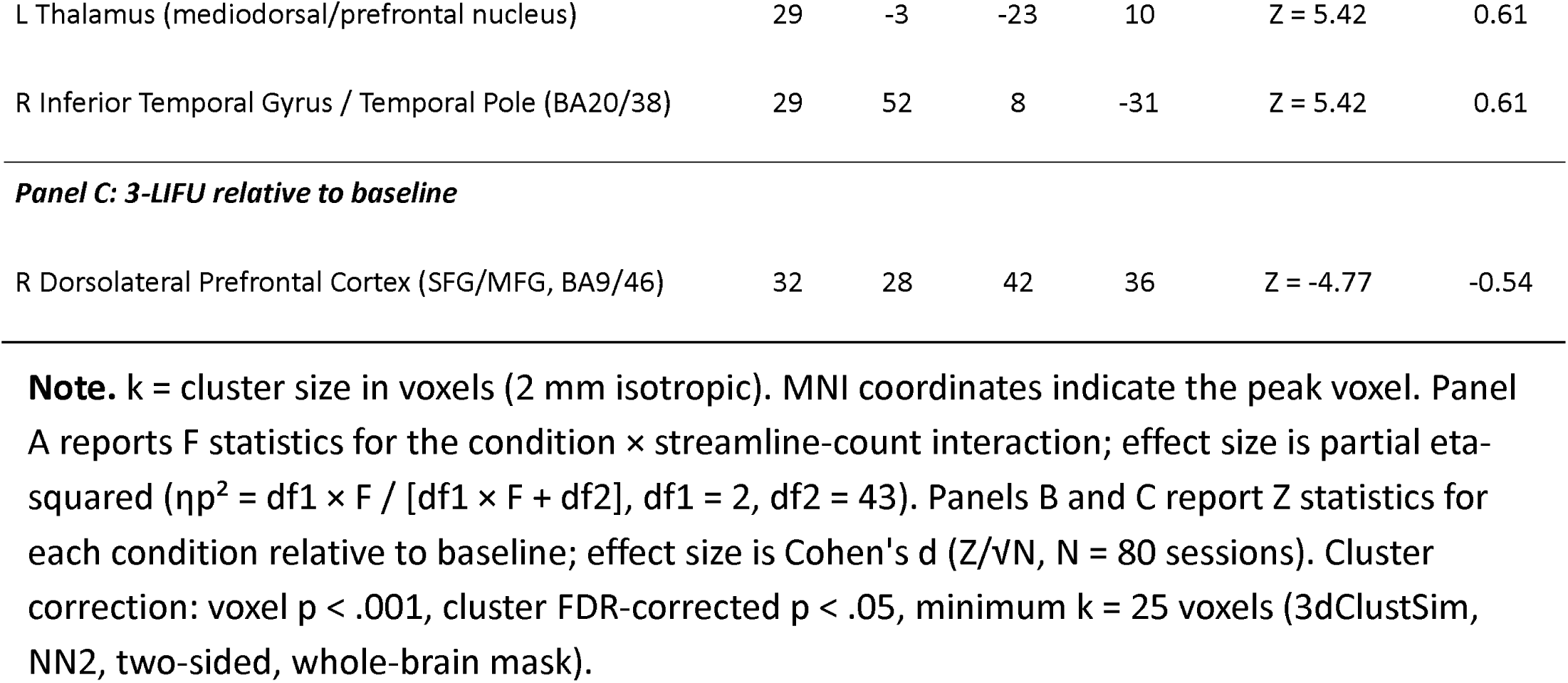
Aim 2: Clusters Associated With Thalamo-Prefrontal Structural Connectivity Moderating LIFU-Induced Functional Connectivity Changes.

### Adverse Events and Tolerability

Most participants reported transient pressure discomfort on the scalp where the ultrasound transducer contacted the skin. The sensation resolved after a few minutes and was considered study-related and expected given that the transducer remained positioned for more than 45 minutes. One participant developed symptoms compatible with a vasovagal episode 15 minutes after 3-LIFU sonication, which resolved within 5 minutes after terminating the scanning session, as monitored by a board-certified emergency medicine specialist (AT). No immediate or delayed pain syndromes, emotional changes, or thermal sensations were reported.

### Blinding

Across 69 sonication visits from 35 participants, participants correctly identified the assigned condition in 33 visits (47.8%; 95% CI, 35.6%-60.2%), which did not differ from chance (binomial p = .810; Bang’s blinding index = -0.04, 95% CI, -0.29 to 0.20). Confidence ratings did not distinguish correct from incorrect guesses (Mann-Whitney U = 581.5, p = .868), and participant guesses were not associated with assigned condition (Fisher exact p = .811; OR = 0.84, 95% CI, 0.29-2.39). Detailed condition- and visit-specific analyses are provided in the Supplement.

## Discussion

The present study describes a preregistered, randomized, double-blind trial of two LIFU doses applied to deep white matter tracts in healthy adults. The main findings were (1) the two doses produced different right-thalamic functional connectivity patterns; (2) three LIFU sonications produced an initial connectivity decrease followed by progressive recovery over the 30-min observation period, with a trajectory that differed from one sonication; and (3) functional connectivity changes were strongly related to the structural connectivity of the targeted white matter pathways, measured as streamline density in the probabilistic tractography model.

Although the present human study cannot resolve the cellular mechanism of LIFU effects in white matter, preclinical evidence provides a plausible framework linking ultrasound, myelinated axons, and circuit-level functional connectivity. Ultrasound can attenuate conduction in myelinated axons (24,28,34–40). Mechanosensitive two-pore-domain potassium channels enriched at nodes of Ranvier, including TRAAK and provide a candidate substrate for this effect, given that the cellular mechanism of repolarization at this level does not align with the classic Hodgkin-Huxley model^41^. Activation of nodal conductance could transiently hyperpolarize or shunt nodes, thereby altering saltatory conduction across fibers traversing the sonication focus. At the mesoscopic level, such a perturbation could initially appear as increased BOLD functional connectivity if simultaneous modulation of connected regions increases temporal covariance. Functional connectivity reflects correlated activity and does not by itself specify excitation, inhibition, or preserved information transfer. With greater or more sustained engagement, the same process could manifest as reduced functional connectivity, followed by recovery as nodal excitability and axonal conduction return toward baseline. This mechanism offers a testable bridge between nodal biophysics in sonicated white matter and the dose- and time-dependent connectivity changes observed here. However, it remains speculative and does not exclude other ultrasound-sensitive processes, including glial effects that may be particularly relevant to white matter, as reviewed elsewhere (42).

To our knowledge, this study provides the first in vivo human evidence of dose-related LIFU effects on deep white matter function. If replicated in larger samples and independent cohorts, the implications may be broad. LIFU of white matter tracts could suppress circuit function and enable causal studies of the neural bases of normal and abnormal behavior. The identification of a LIFU dose that was well tolerated and biologically active in healthy participants may also motivate research into causal circuit-symptom relationships and potential therapeutic applications in clinical samples. Chronic attenuation of axonal transmission in purportedly hyperactive limbic circuits that sustain psychiatric symptoms, including circuits disrupted by capsulotomy, anterior cingulotomy, subcaudate tractotomy, and limbic leukotomy, could, in theory, produce neuroplastic changes and long-term symptom modification (43–47). LIFU could also be used to test the function and effects of deep brain circuits before invasive or irreversible interventions such as ablation or chronic stimulation (48).

The blinding assessment supports the internal validity of the one-epoch versus three-epoch comparison. At the group level, participant guesses were at chance, were not associated with assigned condition, and were not predicted by confidence ratings in mixed-effects models. Four high-confidence correct guesses suggest that occasional partial unblinding may occur through subjective somatic sensations, particularly during higher-dose stimulation, even when overall blinding is preserved. None of these participants attributed the guess to an auditory cue. These findings support the feasibility of blinding in LIFU dose-comparison studies while underscoring the importance of routine post-session blinding assessment, including sensitivity analyses excluding these four visits.

Several limitations should be considered. First, although stimulation targets were defined using individualized tractography and acoustic modeling, the final sonication target remained subject to imprecision, including within-participant variability across the one- and three-epoch conditions. Second, we lacked independent in vivo verification of the actual site and spatial extent of sonication and therefore cannot determine with certainty how closely the modeled target overlapped the effective field in each participant. Third, the 41-min scans raise the possibility that fatigue, motion, physiological drift, or other nonspecific time-dependent effects contributed noise to the functional connectivity estimates. Fourth, the approach used to estimate the direction and depth of the acoustic focus assumed idealized acoustic conditions and did not account for participant-specific variation in cranial shape, thickness, and acoustic properties. These factors may alter the shape and energy of the ultrasound focus and add heterogeneity to the results. Future analyses may partially mitigate these uncertainties by focusing on region-to-region connectivity between gray matter nodes connected most specifically by the sonicated white matter bundles. Nonetheless, even this more anatomically personalized approach would offer only a partial solution because residual variability in tractography-based targeting and transcranial acoustic propagation is expected to remain. Finally, the cerebellar finding lies near the edge of the imaging coverage, making it less reliable.

These limitations notwithstanding, the present study demonstrates dose-dependent functional connectivity responses to two LIFU doses applied to deep white matter tracts in humans. Characterizing these dose-dependent phenomena may be a step toward trials designed to establish causal circuit-symptom relationships and advance circuit-specific therapeutics: reversibly with LIFU or other noninvasive neuromodulation methods, permanently with ablative procedures, including high-intensity focused ultrasound, or chronically with deep brain stimulation.

## Acknowledgments

This work was partly funded by The William K. Warren Foundation, the National Institute of Mental Health (R21 MH138911 and R01 MH137331 to SMG, U01 MH123427 to NSP, and R01 MH132565 to MI), the National Institute of General Medical Sciences Center (Grant 2 P20 GM121312 to MPP), the US Department of Veterans Affairs (I50 RX002864 and I01 RX005299 to NSP), and the National Institute on Drug Abuse (U01 DA050989 to MPP). This paper reflects the opinions of the authors and does not represent the position or policy of the National Institutes of Health or the Department of Veterans Affairs.

