## Supplemental File for "Dose-Dependent Effects of Low-Intensity Focused Ultrasound on Human Deep White Matter Tracts"

**MRI Acquisition**: Resting-state fMRI data were acquired on a GE 3T scanner using a 16-channel head coil apt to accommodate the LIFU transducer. The baseline visit included T1-weighted anatomical imaging, diffusion-weighted imaging, and a 6-min resting-state fMRI scan. Each LIFU visit included a continuous 41-min resting-state fMRI acquisition consisting of a 6-min prestimulation baseline, a 5-min stimulation-delivery period, and a 30-min poststimulation period. The T1-weighted images were acquired using a magnetization-prepared rapid gradient-echo (MPRAGE) sequence with the following parameters: FOV = 256 mm, matrix = 256 × 256, voxel size = 1.0 mm isotropic, 208 sagittal slices, TR/TE = 6/2.92 ms, SENSE acceleration factor R = 2, flip angle = 8°, inversion time (TI) = 1400/1060 ms, sampling bandwidth = 31.25 kHz, scan time = 6 min 11 s. Resting-state fMRI data were acquired using a single-shot gradient-recalled echo-planar imaging (EPI) sequence with sensitivity encoding (SENSE) and the following parameters: TR/TE = 2000/27 ms, flip angle = 78°, FOV/slice = 240/2.9 mm, 39 axial slices, SENSE acceleration factor R = 2, 96 × 96 acquisition matrix reconstructed to 128 × 128, resulting in a voxel size of 1.875 × 1.875 × 2.9 mm³, sampling bandwidth = 250 kHz, scan time = 6 min (180 TRs).

Diffusion-weighted imaging parameters at baseline were as follows: a multiband sequence with an acceleration factor of 3 and 102 diffusion-encoding directions acquired across four shells (b = 500, 1000, 2000, and 3000 s/mm²), TR/TE = 4100/81.7 ms, acquisition and reconstruction matrix = 140 × 140, field of view (FOV) = 24.0 × 24.0 cm, slice thickness = 1.7 mm, and 12 non-diffusion-weighted images. A reverse-phase-encoding acquisition comprising six non-diffusion-weighted images (b = 0 s/mm²) and six diffusion-weighted images (b = 3000 s/mm²) was also obtained to correct for EPI image distortions. The total acquisition time was 7 min 27 s.

**Functional Connectivity Analysis**

BOLD data were preprocessed with fMRIPrep v25.2.5 using MNI152NLin2009cAsym 2-mm standard-space output and a sessionwise anatomical reference. Three initial dummy volumes were discarded. Nuisance regression was then performed in AFNI v25.2.16 using a 12-parameter motion model (six rigid-body motion parameters and their temporal derivatives), five white-matter and five cerebrospinal-fluid aCompCor components, and 13 physiological regressors derived from respiratory and cardiac recordings, consisting of five respiratory-volume-per-time regressors and eight RETROICOR regressors. The time series were bandpass filtered from 0.01 to 0.1 Hz and spatially smoothed within the brain mask using a 4-mm full-width-at-half-maximum kernel. AFNI was also used for temporal filtering, motion censoring, seed-based functional connectivity estimation, and group-level mixed-effects modeling.

Motion-contaminated volumes were censored before functional connectivity estimation. Volumes were censored if framewise displacement exceeded 0.3 mm or if standardized DVARS exceeded 1.5; one additional TR before and one after each flagged volume were also censored. Sessions with more than 50% censored volumes were excluded from analysis. After motion-based exclusions, the functional connectivity analyses included 80 sessions from 31 participants.

Seed-based connectivity was estimated by computing Pearson correlations between the mean time series of the right thalamus seed from the Brainnetome Atlas (1,538 voxels) and the time series of every other brain voxel. Correlation coefficients were Fisher z-transformed before group-level analysis. The 30-min poststimulation period was divided into five consecutive 6-min windows (180 volumes per window). The single 6-min resting-state scan from the no-LIFU baseline visit served as the common reference for each of the five poststimulation windows in both LIFU conditions.

Group-level analyses were conducted in AFNI using linear mixed-effects models (3dLME) with subject-specific random intercepts and Type III sums of squares. Age, sex, and window-matched mean framewise displacement were included as covariates. The preregistered Aim 1 model tested poststimulation functional connectivity as a function of condition (baseline, 1-LIFU, and 3-LIFU). Aim 2 tested whether total thalamo-prefrontal structural connectivity (defined as the sum of probabilistic tractography streamlines connecting the right thalamus with the right anterior cingulate cortex and right orbitofrontal cortex) moderated stimulus-related functional connectivity changes. Aim 3 tested condition-by-time window interactions across the 30-min poststimulation recovery period, including post hoc linear trend and decay contrasts. An exploratory analysis examined within-visit changes in functional connectivity from pre-LIFU to poststimulation for the LIFU visits only.

Whole-brain cluster correction was performed with AFNI 3dClustSim using spatial autocorrelation-function parameters estimated from the residuals (a = 0.611, b = 4.016, c = 2.180; NN2 connectivity; two-sided testing). A voxel-level threshold of p < 0.001 was combined with a cluster-corrected threshold of p < 0.05, corresponding to a minimum cluster extent of 25 voxels within the whole-brain mask (268,341 voxels).

**Blinding Assessment Results**. Confidence ratings did not differ between correct and incorrect guesses (Mann-Whitney U = 581.5, p = 0.868). In a mixed-effects logistic regression that included actual treatment, visit, confidence rating, and a random participant effect, actual treatment (p = 0.790), visit (p = 0.148), and confidence rating (p = 0.472) did not predict correct identification. Of 69 free-text responses, 20 (29%) mentioned a physical sensation and 52 (75%) described the response as a pure guess; mentioning a physical sensation was not significantly associated with correct identification (p = 0.110; OR = 2.65, 95% CI, 0.81–9.37). Four participants made a high-confidence correct guess on one visit each; high confidence was prespecified as a rating ≥ 6 on a 0–10 scale. Although all four high-confidence guesses were correct, this subgroup was small and did not differ significantly from chance (4/4 correct; p = 0.125). Three high-confidence correct guesses occurred after three-epoch LIFU and were attributed by participants to subjective head sensations, pulses, or surges. One occurred after one-epoch LIFU and was attributed to the absence of any perceived difference. None was attributed to an auditory cue. Overall, these findings support successful blinding at the group level while identifying rare sessions in which subjective somatic sensations may have contributed to partial unblinding.

**Supplementary Table 1. Inclusion and exclusion criteria.**

| **Category** | **Domain** | **Criterion** |
| --- | --- | --- |
| **Inclusion** | Age | 18 to 65 years. |
| **Inclusion** | Language and consent | English fluency and capacity to provide written informed consent. |
| **Inclusion** | Psychiatric status | No lifetime DSM-5-TR psychiatric diagnosis, as confirmed by the MINI structured interview and consultation with a board-certified psychiatrist. |
| **Inclusion** | Pregnancy | For participants of childbearing potential, a negative urine pregnancy test at screening. |
| **Inclusion** | Incidental findings | Consent to the possibility of incidental findings, such as a brain abnormality detected during imaging. |
| **Exclusion** | Suicide risk | Active suicidal ideation on C-SSRS, defined as yes responses to items 3, 4, or 5 in the past-month suicidal ideation section, any yes response to suicidal behavior in the past 3 months, or any suicide attempt in the past 3 months. |
| **Exclusion** | Substance or alcohol use disorder | History of moderate or severe substance or alcohol use disorder according to DSM-5-TR; a substance use disorder of greater than mild severity within the past 6 months was also exclusionary. |
| **Exclusion** | Alcohol or drug screen | Positive screening result for alcohol or drugs of abuse, including methadone, opiates, cocaine, amphetamine/methamphetamine, ecstasy, or other prohibited substances. |
| **Exclusion** | Psychiatric diagnosis | Current or lifetime DSM-5-TR psychiatric disorder. |
| **Exclusion** | Concomitant medications | Use of benzodiazepines or anticonvulsants in the 7 days before screening, or prescription medication outside the accepted range as determined by clinical best practice and current research. |
| **Exclusion** | MRI feasibility or safety | Inability to complete MRI scanning or presence of an MRI contraindication identified during safety screening. |
| **Exclusion** | Central nervous system pathology | Clinical history of relevant central nervous system structural pathology, including Parkinson disease, multiple sclerosis, or malignant brain neoplasm. |
| **Exclusion** | Medical stability | Uncontrolled diabetes mellitus, defined as fasting glucose ≥ 120 mg/dL or hemoglobin A1c ≥ 6.5%, or uncontrolled hypertension, defined as two consecutive readings ≥ 140/90 mmHg. |
| **Exclusion** | Cognition | Clinical history of at least minor neurocognitive disorder of any origin. |
| **Exclusion** | Capacity or protocol adherence | Medical, psychiatric, or other condition limiting the ability to understand study information, provide informed consent, adhere to protocol requirements, or complete the study. |
| **Exclusion** | Communication | No reliable means of communication, such as lack of internet or telephone access. |
| **Exclusion** | Protocol completion | Unwillingness or inability to complete any major aspect of the study protocol. |
| **Exclusion** | Sensory function | Uncorrectable visual or hearing impairment. |

Abbreviations: C-SSRS, Columbia-Suicide Severity Rating Scale; DSM-5-TR, Diagnostic and Statistical Manual of Mental Disorders, Fifth Edition, Text Revision; LIFU, low-intensity focused ultrasound; MINI, Mini International Neuropsychiatric Interview; MRI, magnetic resonance imaging.

**Supplementary Table 2. Assessment of participant blinding for one-epoch and three-epoch LIFU**

Summary of forced-choice postscan guesses, confidence ratings, formal blinding tests, and interpretation of results.

| **Domain** | **Metric** | **Result** | **Interpretation** |
| --- | --- | --- | --- |
| Analysis sample | 69 stimulation visits; 35 participants | 69 completed exit-questionnaire guesses; assignment and guess available for all 69 visits. | Complete blinding dataset available for the current analysis. |
| Overall guessing accuracy | 33/69 correct | 47.8% correct; 95% CI, 35.6%-60.2%; binomial p = 0.810; Bang's BI = -0.04 (95% CI, -0.29 to 0.20). | Accuracy did not differ from chance, supporting successful blinding at the group level. |
| Accuracy by actual condition | Actual 1-LIFU: 16/34 correct; actual 3-LIFU: 17/35 correct | 1-LIFU: 47.1% correct, p = 0.864, Bang's BI = -0.06. 3-LIFU: 48.6% correct, p = 1.000, Bang's BI = -0.03. | No evidence that either dose condition was more readily identifiable. |
| Accuracy by visit | Visit 1: 19/35 correct; Visit 2: 14/34 correct | Visit 1: 54.3% correct, p = 0.736, Bang's BI = 0.09. Visit 2: 41.2% correct, p = 0.392, Bang's BI = -0.18. | No evidence that blinding differed reliably by session. |
| Actual assignment versus participant guess | 2 × 2 Fisher's exact test | Actual 1-LIFU: 16 guessed 1-LIFU, 18 guessed 3-LIFU. Actual 3-LIFU: 18 guessed 1-LIFU, 17 guessed 3-LIFU. Overall p = 0.811; OR = 0.84 (95% CI, 0.29-2.39). | Participant guesses were not associated with actual treatment assignment. |
| Confidence versus accuracy | Correct guesses n = 33; incorrect guesses n = 36 | Median confidence was 0.0 for both correct and incorrect guesses; Mann-Whitney U = 581.5, p = 0.868. | Confidence ratings did not distinguish correct from incorrect guesses. |
| High-confidence subgroup | Prespecified confidence ≥ 6 on a 0–10 scale: 4 visits | 4/4 high-confidence guesses were correct; binomial p = 0.125; Bang's BI = 1.00 (95% CI, -0.20 to 1.00). | This qualitative signal was noteworthy, but the subgroup was very small and did not differ statistically from chance. |
| Mixed-effects logistic regression | correct ~ actual treatment + visit + confidence + (1 \| participant) | No significant fixed effects: actual treatment p = 0.790; visit p = 0.148; confidence p = 0.472. Predicted P(correct): 1-LIFU = 0.425, 3-LIFU = 0.470. | Findings remained consistent after accounting for repeated observations within participants. |
| Free-text reasoning | 69 responses | 20/69 (29%) mentioned a physical sensation; 52/69 (75%) described the response as a pure guess. Sensation mention versus correctness: p = 0.110; OR = 2.65 (95% CI, 0.81-9.37). | Subjective sensations were reported by a minority of participants but were not significantly associated with correct identification. |
| High-confidence correct guesses: qualitative review | 4 visits; 4 participants | Three high-confidence correct guesses occurred after 3-LIFU; participants attributed them to subjective head sensations, pulses, or surges. One high-confidence correct guess occurred after 1-LIFU; the participant attributed it to the absence of any perceived difference. None was attributed to an auditory cue. | These rare sessions may represent partial unblinding and should be reported transparently; sensitivity analyses excluding these four visits may be considered. |

Abbreviations: BI, Bang's blinding index; CI, confidence interval; LIFU, low-intensity focused ultrasound; OR, odds ratio. Bang's BI = 2 × proportion correct - 1 in this forced-choice design; 0 indicates chance-level guessing and 1 indicates complete unblinding. Confidence ≥ 6 was prespecified as high confidence.

**Participant Flow Diagram**

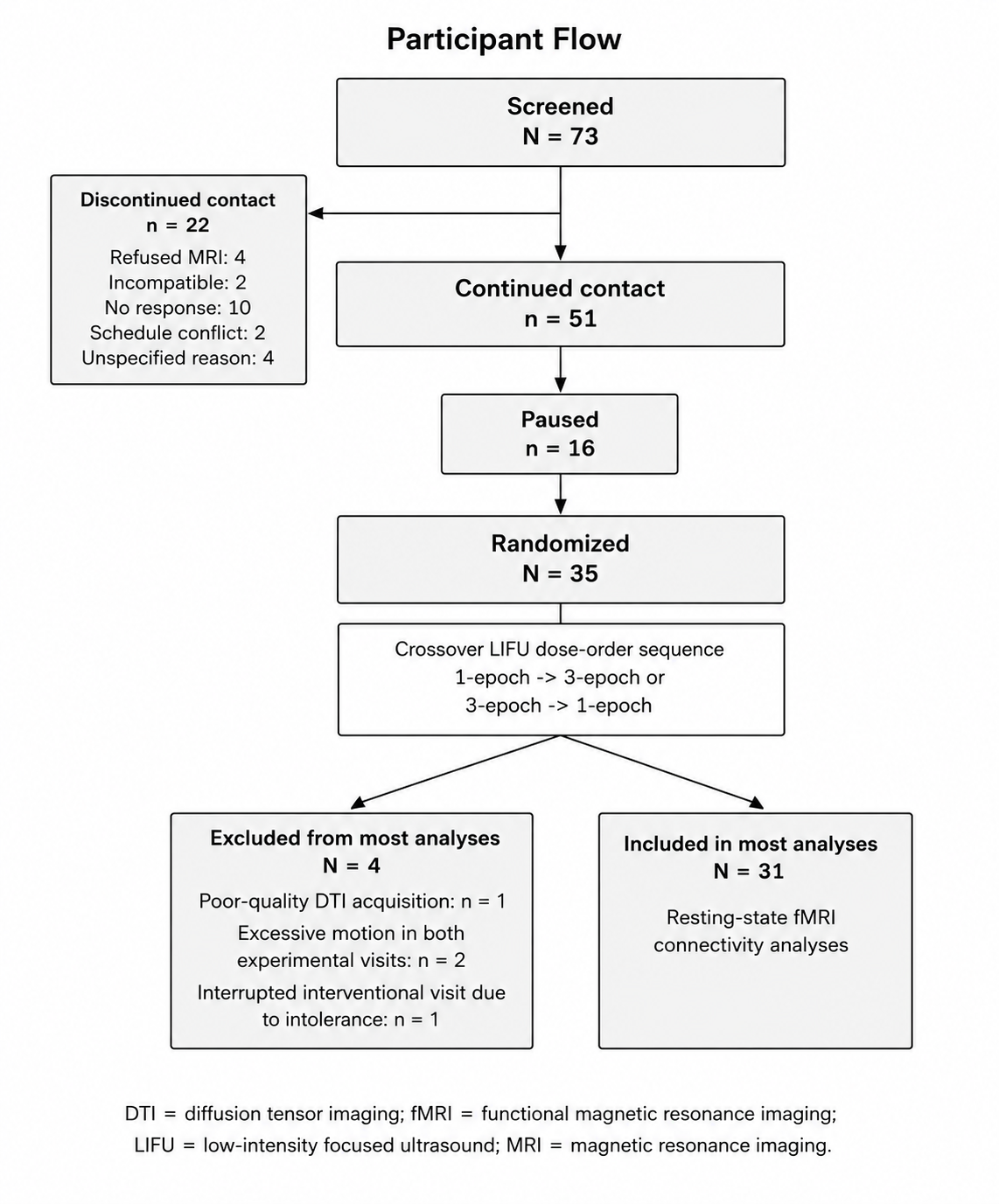
